# Identification of disease mechanisms and novel disease genes using clinical concept embeddings learned from massive amounts of biomedical data

**DOI:** 10.1101/2023.04.27.538319

**Authors:** Andrej Bugrim

## Abstract

**Motivation:** Knowledge of relationship and similarity among human diseases can be leveraged in many biomedical applications such as drug repositioning, biomarker discovery, differential diagnostics, and understanding of disease mechanisms. Recently developed *cui2vec* resource provides embeddings of approximately 109 thousand biomedical terms and allows computing novel measures of disease similarity, directly related to patterns in real-world data. We investigate whether disease embeddings from *cui2vec* can be utilized to identify functional relations among diseases, to uncover their molecular mechanisms, and to generate hypotheses about novel gene-disease associations and potential drug targets.

**Methods and results:** We focus on a subset of 3,568 *cui2vec* terms corresponding to human diseases annotated in DisGeNET database. Disease-disease distance matrix is computed for this set of diseases based on their embedding vectors. Clustering of this matrix reveals a well-defined structure with good correspondence between disease clusters and the top MeSH disease categories. Using pulmonary embolism as an example we show how disease clustering is related to known mechanistic relations among diseases. Next, we combine disease embeddings with annotated gene-disease associations from DisGeNET to generate joint gene-disease co-embeddings. From these we identify molecular pathways most characteristic for each disease group and show that they are highly relevant to known disease physiology. Finally, we leverage disease similarity to generate and rank hypothesis for gene-disease associations and demonstrate that this method generates highly accurate results and can suggest relevant drug targets.

**Conclusions:** We show that combination of disease embeddings learned from massive amounts of biomedical records with curated data on gene-disease associations can reliably reveal groups of functionally related diseases and their molecular mechanisms and predict novel gene-disease associations. Importantly, our analysis does not require knowledge of associated genes for every disease to identify patterns in the embedding space, therefore it can be used to suggest mechanisms for conditions that have not been functionally understood. In this respect our analysis can be applied to identify potential markers and drug targets for poorly characterized orphan and rare diseases. It can also reveal unexpected novel connections among diseases and between diseases and molecular pathways.

## Introduction

Knowledge of relationship and similarity among human diseases has been leveraged for understanding molecular disease mechanisms (1-4), repositioning drugs (5-7), aiding differential diagnosis of poorly understood conditions (8) and other important tasks in research, clinical practice, and drug development. Until recently the most common ways to define disease similarity were either based on disease taxonomy (9-11), symptoms or phenotypic similarity (12-15), shared genes and/or molecular mechanisms (16,17) or combinations of these factors (18). While useful in many applications, these approaches are limited to conditions with well-understood phenotypes and etiology and are based on the existing knowledge which is incomplete and often biased. They are not very conducive for the discovery of novel relations among pathologies and cannot handle poorly understood conditions, notably many rare diseases.

With the development of large electronic repositories of biomedical information, it became possible to analyze massive amounts of Electronic Health Records (EHRs), insurance claims, medical notes, biomedical literature, and other documents to generate the so-called *vector embeddings* of biomedical concepts. This approach is based on word embedding models of natural languages. In this method, each term/word is represented as a vector of real numbers in the embedding space with the goal that similar and related terms are placed close to each other. There are several popular algorithms for generating word embeddings from massive amounts of text documents, including word2vec (19), GloVe(20), and FastText (21). When applied to the analysis of health-related and biomedical documents these and related methods can generate representations of biomedical terms including human diseases (22-27). Conditions that often co-occur in the same patient or are often mentioned together in research papers are expected to be placed close to each other in the embedding space, allowing to define a new measure of disease similarity (28). This similarity measure is attractive because it is based on real-world data, does not use any preconceptions about disease mechanisms or relationship among diseases and can include conditions that have not been previously classified or understood mechanistically. Thus, it could allow discovery of novel relations among diseases, can suggest mechanisms for poorly understood pathologies and enable identification of new links between diseases and other concepts, such as drugs and genes.

In this work we leverage the recently developed *cui2vec* resource which provides embeddings of approximately 109 thousand biomedical terms in a 500-dimensional vector space (25). These embeddings are learned from term co-occurrence matrix derived from the combined corpus of 60 million insurance claims, 20 million clinical notes and 1.7 full text papers from PubMed Central. In comparison to other publicly available embedding models of biomedical concepts *cui2vec* is based on the significantly larger document corpus in which data from different sources are combined to generate the single embedding model. According to the authors’ of (25), *cui2vec* outperforms other models in the task of recognition of known relations among biomedical concepts. Here our goal is to investigate whether *cui2vec* data could be used to identify functionally related diseases, to uncover common disease mechanisms, and to generate credible hypotheses about novel gene-disease associations. To this end we combine *cui2vec* data with gene-disease associations obtained from DisGeNET database (29) to generate joint gene-disease embeddings which we leverage for performing functional analysis tasks outlined above.

## Results

### Data overlap between cui2vec and DisGeNET

*Cui2vec* contains approximately 109K biomedical terms of different nature identified by their Concept Unique Identifiers (CUI). DisGeNET contains curated gene-disease associations describing 84,038 links between 9,703 genes and 11,181 pathological conditions. The mapping of CUI identifiers between *cui2vec* and DisGeNET data tables yields 3,568 common concepts, representing ∼32% of all diseases annotated to genes in DisGeNET (Table S1). These diseases participate in 53,032 associations with 8,686 genes. While the overlap represents *approximately* one third of all disease terms in DisGeNET, the majority (∼63%) of gene-disease associations is contained in this set.

### Disease categories emerge from data

In this study, we focus on 3,568 *cui2vec* terms mapped to human diseases annotated in DisGeNET. Disease-disease cosine similarity matrix and corresponding distance matrix are computed for this set of diseases based on their embedding vectors (see *Methods*). Hierarchical clustering reveals a well-defined cluster structure (Fig. 1, Table S1). To verify that embedding-based clustering is functionally meaningful, we map diseases onto 26 top MeSH disease categories (30). Clusters are represented by 26-component vectors with elements corresponding to fractions of clusters’ diseases belonging to each of the MeSH disease types. We observe that each disease cluster is dominated by 1-3 MeSH categories (Fig. 2, Table S2). If multiple categories are represented in the same cluster, they often correspond to related disease types. For example, cluster # 16 is dominated by three MeSH categories: *Mental Disorders, Behavior and Behavior Mechanisms, and Chemically Induced Disorders*.

**Figure 1.**
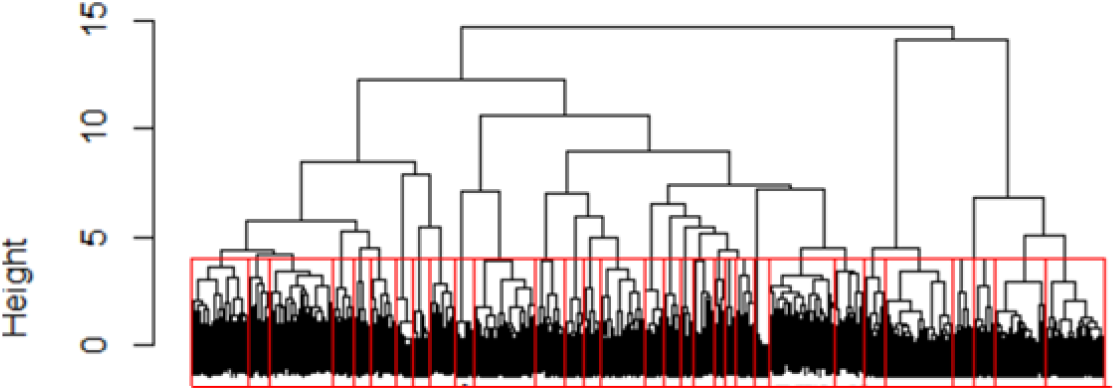
Clustering of 3,568 human diseases in the embedding space based on cosine similarity of their embedding vectors. Red border identifies optimal clusters (described in text)

**Figure 2.**
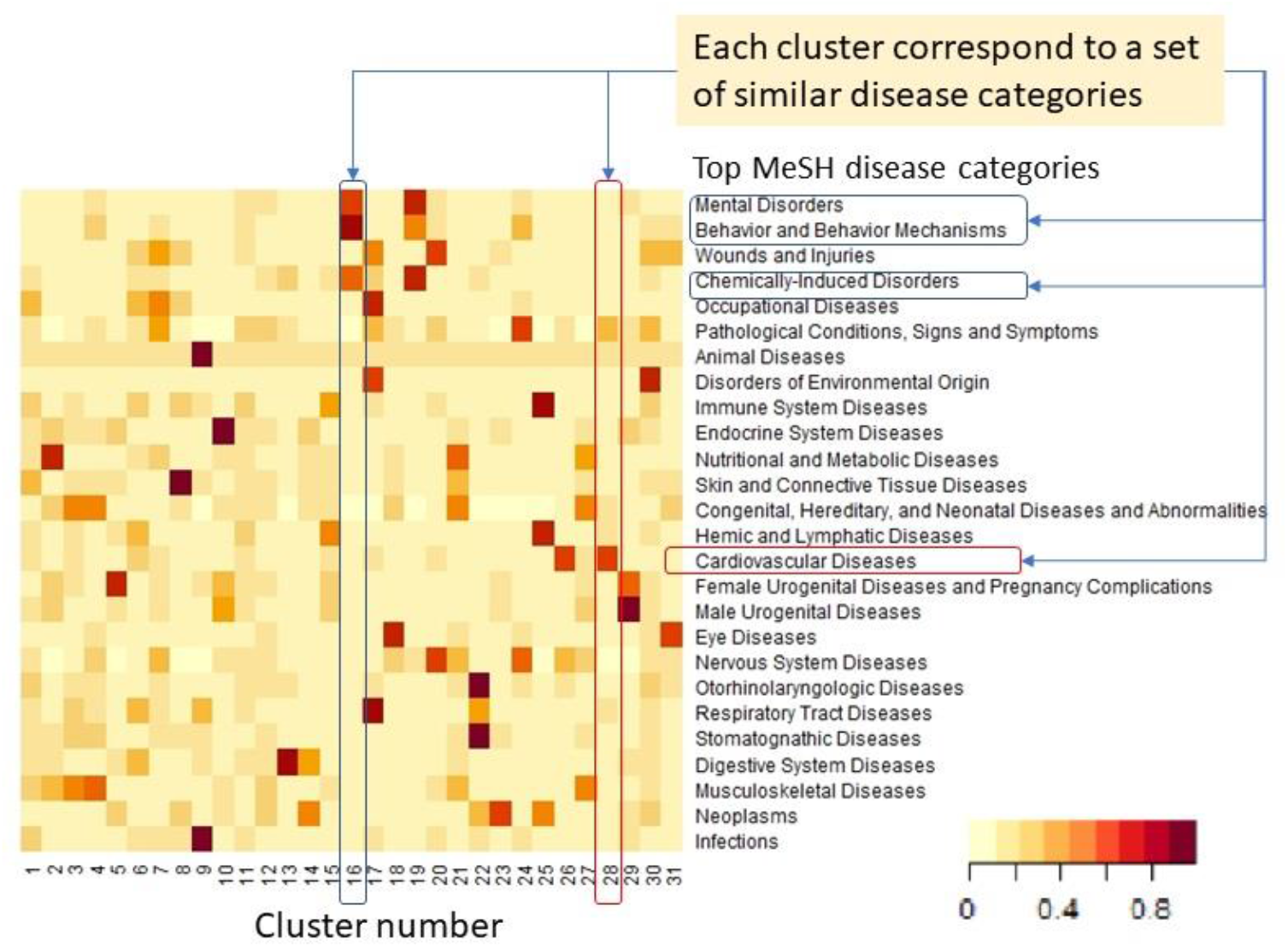
Mapping between disease clusters shown on Fig. 1 and top disease categories from MeSH. Each cluster is dominated by 1 to 3 MeSH categories. Where multiple MeSH categories are represented within the same cluster, they are typically related to each other. Color indicates the fraction of cluster’s diseases annotated to a disease category.

To identify the optimal number of clusters, we attempt to split the set of all diseases into groups with the most functional difference from each other. By progressively decreasing cluster height cut-off we generate increasingly fine-grained sets of disease clusters and compute average pairwise Pearson correlation among columns of the *ClusterXMeSH* matrix depicted on Fig 2. As the number of clusters increases, the correlation drops sharply, indicating that clusters become more functionally independent (Fig. 3). The inflection point is seen as the number of clusters reaches approximately 30. Further increasing the number of clusters does not make them significantly more independent. This leads to the conclusion that the optimal number of clusters is around 30. Incidentally, this number is close to the number of top MeSH categories, which is 26. Together the two results indicate that disease types naturally emerge from the data structure in the embedding space without any prior preconceptions.

**Figure 3.**
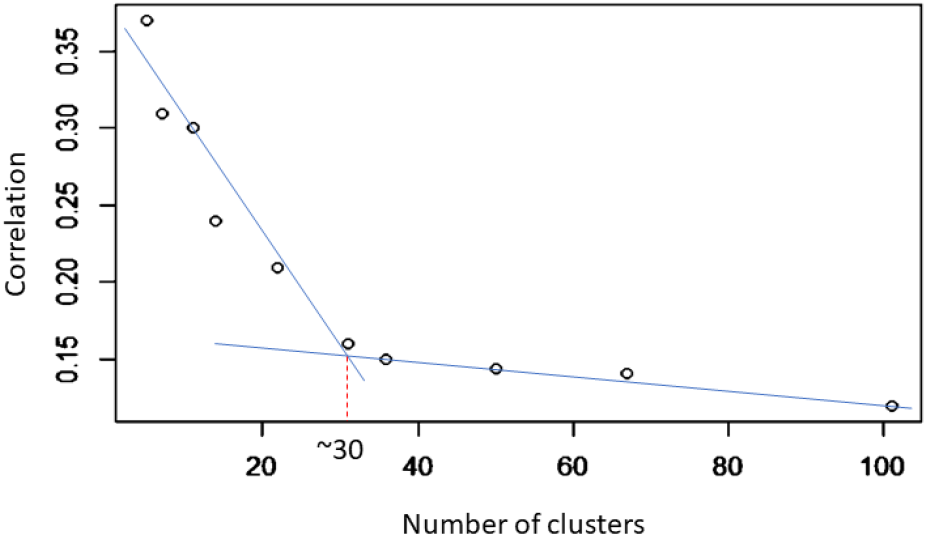
Average correlation between enrichment of different clusters by top MeSH disease categories. The inflection point around 30 clusters indicates the optimal split of diseases into functionally distinct groups.

### Mechanistic basis of disease clustering

To better understand physiological factors that drive disease clustering we perform detailed analysis of pulmonary embolism (PE) and diseases that are close to it in the embedding space. Fig. 4 shows mutual similarity matrix for these diseases with a hierarchy of clusters. At the top level there are two major clusters. Cluster “A” is predominantly comprised by diseases that can be broadly described as conditions associated with or leading to abnormal blood flow. These range from molecular deficiencies directly associated with thrombi formation and different types of thrombosis to embolism and aneurisms. Cluster “A” is further split into smaller, more tightly connected clusters “C”, ”E”, and “D”. Cluster “C” includes different types of vein thrombosis as well as molecular deficiencies that make a person more likely to develop thrombi. Cluster “D” contains aortic aneurisms, another major type of vascular pathology and a cause of abnormal blood flow. It also contains a few general terms like “Thrombosis”, “Thrombus”, and “Embolus” and includes pulmonary embolism, suggesting a clinical connection between aortic aneurisms and thrombi-related diseases. Finally, a small cluster “E” consists of coronary thrombosis, coronary syndrome, and myocardial infarctions. The second top level cluster (cluster “B”) consists of heart conditions and pathologies, some of which may arise as downstream physiological effects of the abnormal blood flow. Interestingly, myocardial infarction clusters with coronary thrombosis rather than with other heart conditions in cluster “B”, probably due to the very high level of comorbidity and well-established clinical connection (31).

**Figure 4.**
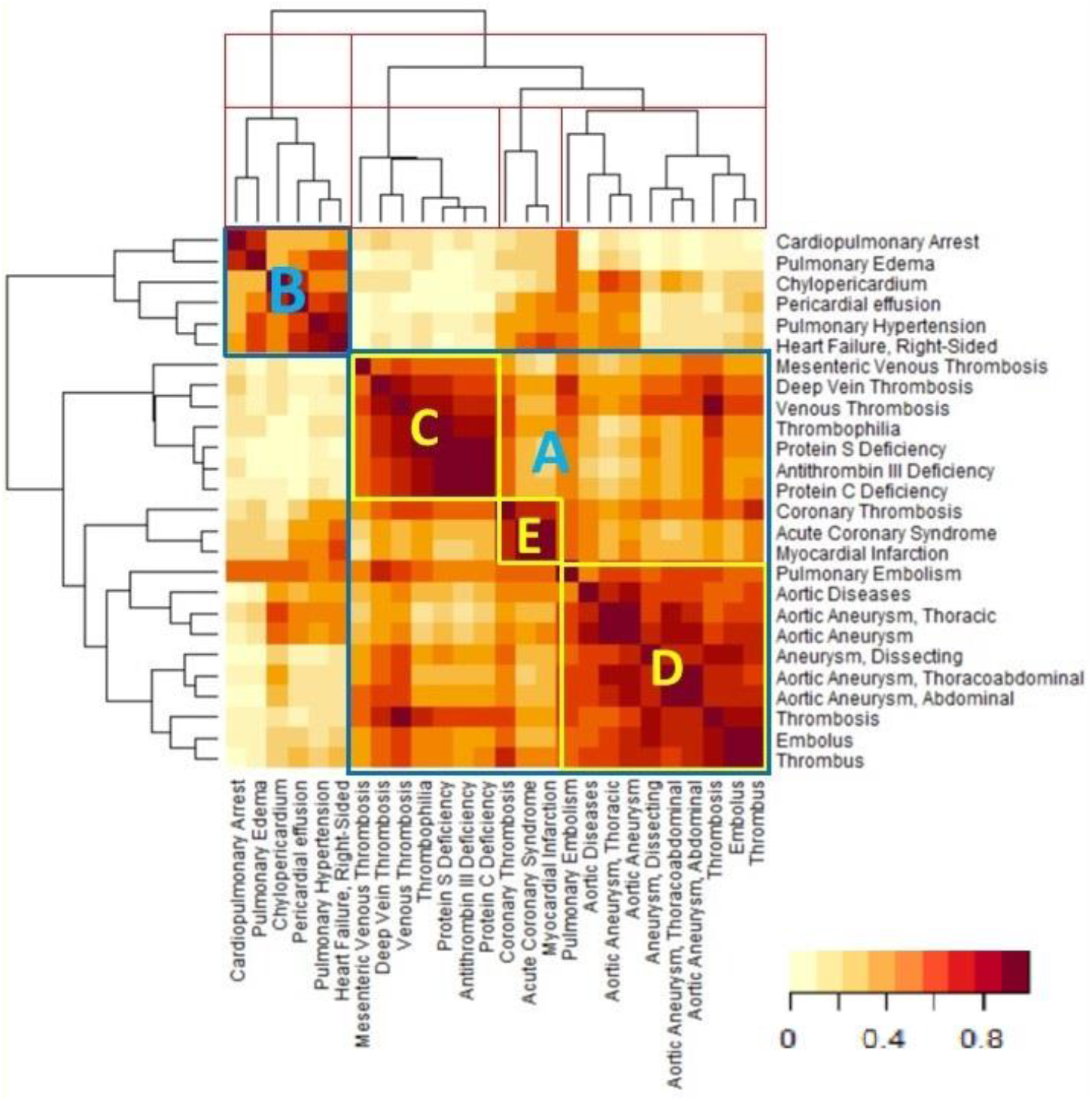
Matrix of the mutual cosine similarity for diseases that are neighbors of pulmonary embolism in the embedding space.

### Gene embedding and functional characterization of disease clusters

The product of gene-disease association matrix and normalized disease embedding matrix yields embeddings of 8,686 genes into the same 500-dimensional vector space in which 3,568 gene-associated diseases are embedded (See *Methods* section for detail). We use this co-embedding to elucidate the most relevant common genes and pathways underlying disease clusters. First, we define embedding vectors for disease clusters as averages of normalized embedding vectors of diseases in each cluster. Second, we compute cosine similarity between clusters’ and genes’ embedding vectors. We consider a gene to be associated with the cluster for which it has the greatest value of cosine similarity. Thus, each gene is associated with one and only one disease cluster. As a result, we obtained 31 non-overlapping gene sets associated with disease clusters. Finally, we compute functional enrichment of these sets using human pathways from Reactome database (32). We find that enriched pathways are highly relevant to the types of diseases represented in each cluster. For example, top pathways for genes associated with cluster #28, consisting of cardiovascular diseases are all associated with the electrophysiology of muscle contraction (Fig. 5A-B). For comparison, we also analyze the set of all genes directly annotated to diseases in this cluster. Pathway enrichment for this set of genes is dominated by inflammation and signal transduction pathways that are much less specific to cardiovascular diseases (Fig. 5C).

**Fig. 5.**
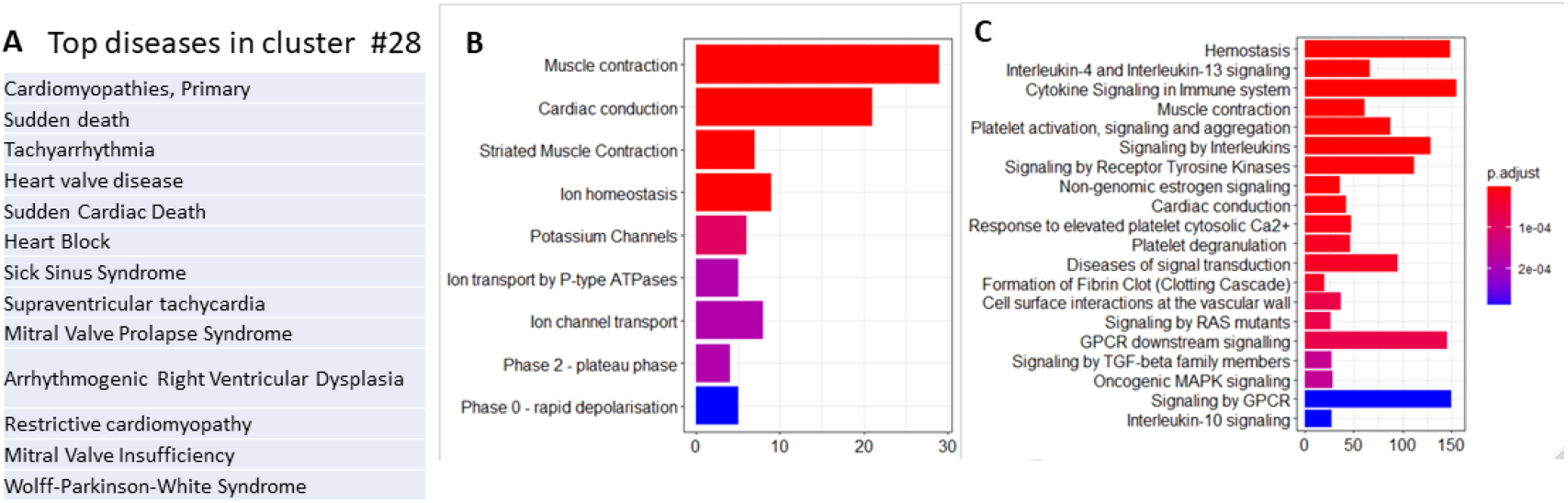
Functional characterization of the cluster of cardiovascular diseases (cluster # 28 on Fig. 2) (A) most diseases in the cluster are related to cardiovascular conditions; (B) top pathways for the set of genes associated with the cluster based on embeddings; (C) top pathways for the union of all genes annotated in DisGeNET database to diseases in the cluster.

To identify the role of disease clustering in the alignment between gene and disease embedding vectors we compute numbers of gene-disease pairs exceeding set similarity thresholds and calculate ratios of the numbers of such pairs obtained with real disease embedding data and those generated using disease embedding matrix with randomized row labels (see *Methods* section). The randomization preserves gene-disease associations but breaks patterns of clustering of diseases annotated to the same gene. Results (Fig. 6.) show that for similarity thresholds above 0.4 the ratio grows sharply, reaching the 7-fold increase for thresholds above 0.8.

**Figure 6.**
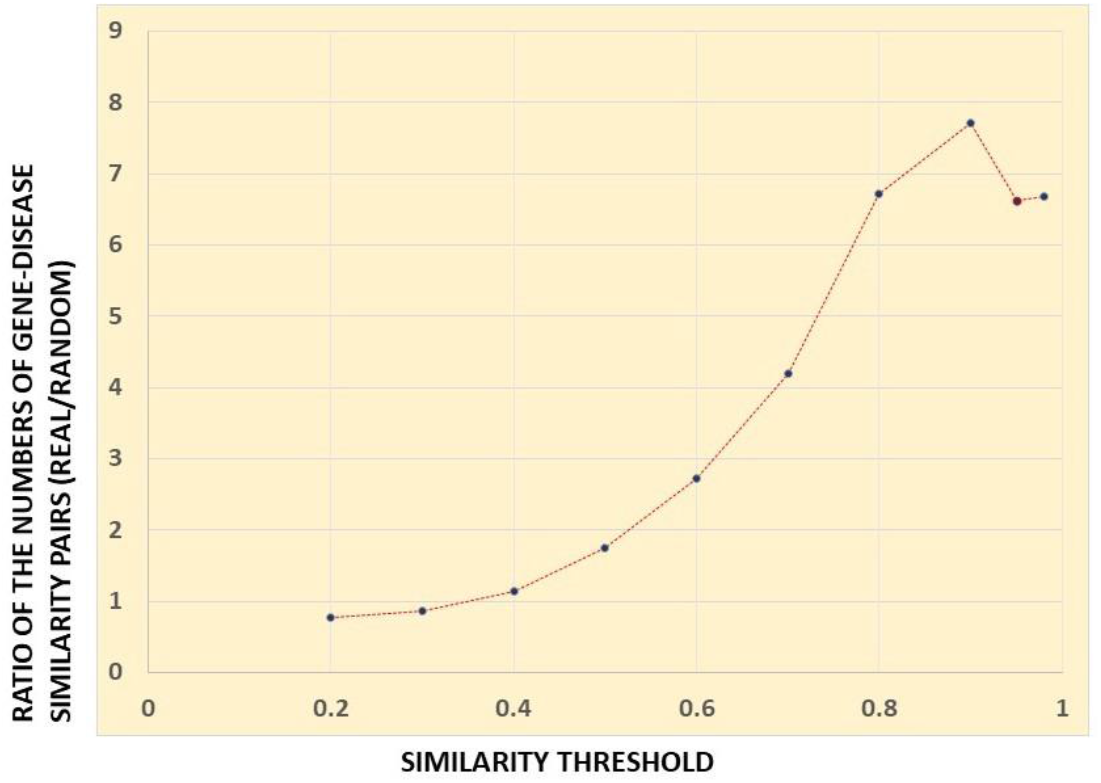
The ratio of numbers of gene-disease pairs exceeding certain similarity thresholds obtained with real and randomized disease embeddings.

The most likely explanation for this result is that diseases associated with the same gene tend to be aligned to each other. This would lead to gene embedding vectors being aligned with multiple disease vectors, resulting in a significant number of high-similarity gene-disease pairs in real data. To test this hypothesis, we compute average similarity within sets of diseases associated with individual genes. The analysis is limited to 3,063 disease sets corresponding to genes with at least 5 associated diseases. The average within-set cosine similarity of diseases is 0.26 compared to 0.09 average similarity among all 3,325 diseases comprising these sets. Notably, the highest within-set disease similarity is observed for sets of a relatively small size. For example, for sets containing 10 or fewer diseases associated with the same gene the average value of similarity is 0.28, whereas for sets containing more than 50 diseases this value drops to 0.16. These results support our hypothesis that clustering of diseases significantly affects the alignment between gene and disease embedding vectors and plays a crucial role in functional characterization of disease clusters.

### Generating hypotheses about novel gene-disease associations

The results described above demonstrate that diseases tend to form functionally meaningful clusters in the embedding space, that this clustering can be attributed to common molecular mechanisms, and that disease embedding vectors are often aligned with those of the genes that are mechanistically relevant to their etiology. We leverage these findings to generate hypotheses about novel disease-related genes using known gene-disease associations and disease similarity. We speculate that a known gene-disease association serves as partial evidence for that gene’s associations with similar diseases. To evaluate the likelihood of a putative gene-disease association we compute *aggregated evidence score*, integrating partial evidence from multiple similar diseases. In computing this score, we emphasize contributions from highly similar diseases, while downgrading those from the multitude of weakly similar ones. This is achieved by using transformed disease similarity matrix obtained by applying S-shaped transforming function that reduces contributions from diseases with cosine similarity < 0.8 (See *Methods* section for detailed description). We compute aggregated evidence scores for all pairs between 3,568 diseases and 8,686 genes in our data set. Using score values as classifiers of gene-disease associations we calculate areas under the ROC curves (AUC) for their ability to differentiate between associated and non-associated genes for each disease. To generate meaningful ROCs this analysis is limited to diseases with at least 5 associated genes in DisGeNET data (1,473 diseases). The results show that aggregated evidence scores are strongly predictive of gene-disease associations. The average value of AUC across all analyzed diseases is 0.89, while for 881 diseases it exceeds 0.9.

To illustrate the mechanistic basis for this method we return to the example of pulmonary embolism (PE) described earlier. DisGeNET contains 12 genes associated with this disease. We identified 30 additional genes for which aggregated evidence score is > 1 and which we label as “probable PE genes”. Figure 7 illustrates different patterns in which their associations with other diseases contribute to the aggregated evidence score for their putative association with PE. Firstly, there is a group of genes annotated to deep vein thrombosis (F5, F2, F3, SERPINC1 and a few others). The mechanistic connection of these genes to pulmonary embolism is straightforward. Blood clots forming in deep veins and then traveling to lung is the primary cause of this disease. Thus, genes that are involved in thrombi formation are also likely to be related to the risk of pulmonary embolism, even though they are not directly annotated to it in DisGeNET. The second group contains genes annotated to diseases that are not directly related to PE or thrombosis. This group consists of MMP2, MMP9, TGFBR2, and ACTA2 which are annotated to different types of aneurisms, myocardial infarction, and heart failure. All these genes play a role in smooth muscle cells migration and vascular remodeling (33-35). Their likely mechanistic connection to PE is mediation of its consequences: vasculopathy following development of chronic pulmonary hypertension and right ventricular failure (36).

**Figure 7.**
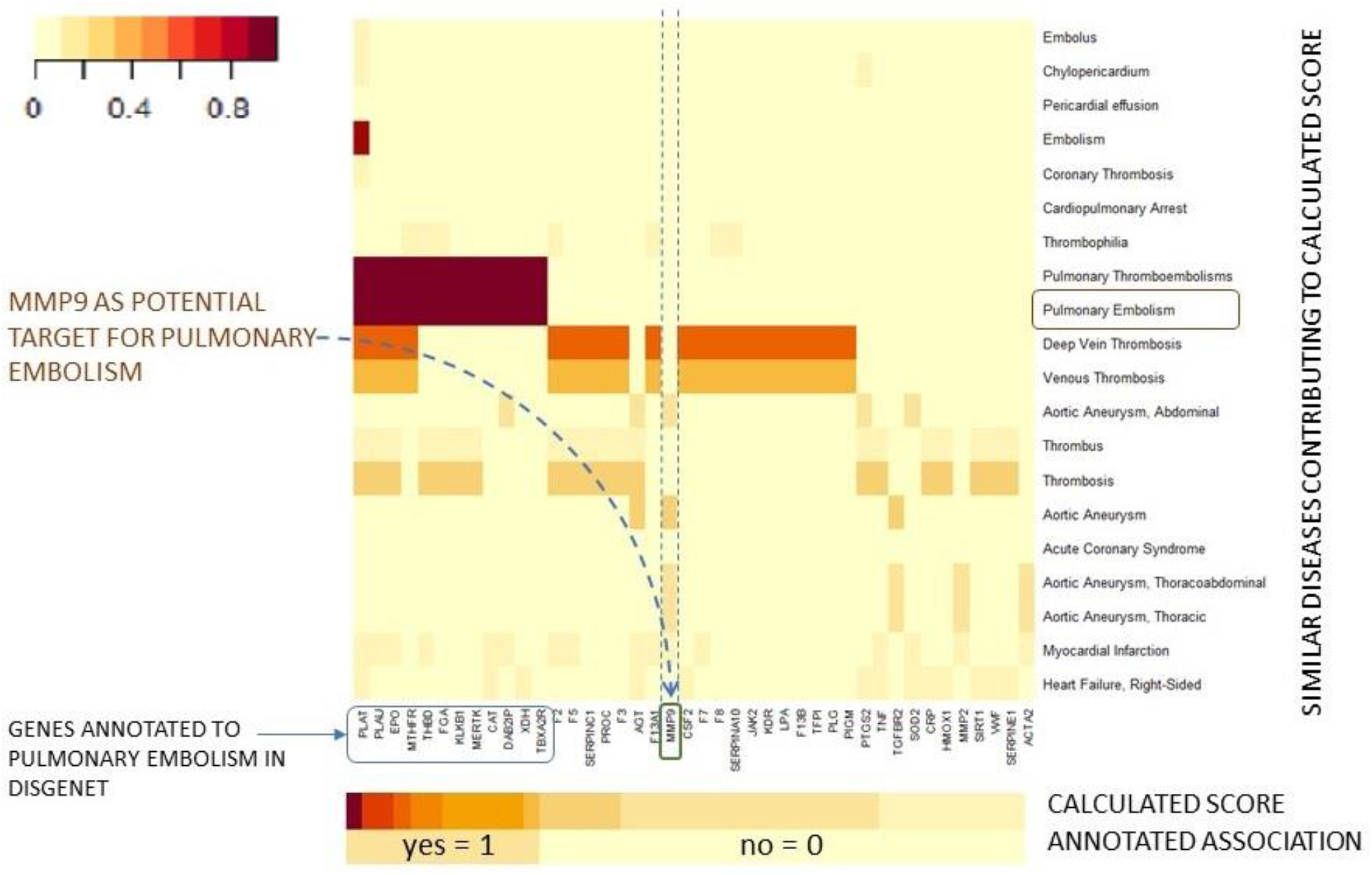
Contribution of different diseases to aggregated evidence scores for established and predicted gene-disease associations of pulmonary embolism.

## Discussion

In this work we analyze human disease embeddings from *cui2vec* learned from medical term co-occurrence statistics in the EHR and other relevant medical and biomedical documents (25). Even though no knowledge about disease mechanisms or disease classification was utilized by the creators of *cui2vec* in computing these embeddings, we find that clustering of diseases based on cosine similarity of their embedding vectors is closely aligned with expert-designated disease categories. Thus, it can be said that groups of functionally related diseases “naturally emerge” from patterns in disease embedding data. Detailed investigation of disease cluster around pulmonary embolism reveals that disease grouping in the embedding space can be attributed either to commonality of underlying mechanisms across different pathologies or to physiological hierarchy with one condition being a cause of another. For example, molecular deficiencies, such as Protein C and Protein S deficiency make a person more likely to develop thrombi. Incidentally, they are a part of a tightly connected group of diseases that includes different types of vein thrombosis (cluster “C” on Fig 4). Diseases in this cluster also share a high degree of similarity with pulmonary embolism which often results from thrombi traveling from deep vein to lung. The most likely reason that these conditions cluster together in the embedding space is their comorbidity in patients who are predisposed to thrombi formation, leading to frequent co-occurrence in medical records. A separate group of closely related diseases predominantly consists of different types of aortic aneurysms (cluster “D” on Fig. 4). Explaining their similarity to pulmonary embolism is not as straightforward as that for thrombosis. The evidence for comorbidity of aortic aneurisms and pulmonary embolism does exist in the literature, but it is limited (37-40). At the same time, embedding patterns suggest that in real life the clinical connection between these conditions is strong. This implies the existence of an unidentified physiological or molecular link. One hypothesis that can be put forward is that high level of activity of genes involved in vascular remodeling (such as matrix metalloproteinases MMP9, MMP2) makes a person predisposed to both: developing aortic aneurisms and suffering more severe consequences as a result of pulmonary embolism. While the role of these proteins in aortic aneurisms is well documented (41,42), the evidence has also emerged that inhibition of MMP9 in laboratory animals helps to alleviate effects of pulmonary embolism (43-46) providing some support for this hypothesis. Another group of diseases (cluster “B” on Fig. 4) is a set of heart conditions. Some of these, such as heart failure and cardiac arrest are downstream effects of PE and are immediate causes of mortality in PE patients. It is interesting to note that while these conditions are aligned to PE in the embedding space, overall, there is not much similarity between diseases in cluster “B” and those in cluster “C”. This may reflect the fact that in general, heart conditions are attributed to many different causes, while at the same time thrombi formation leads to many problems with PE and subsequent heart failure being only one of its consequences. This can be contrasted with the observation that myocardial infarction and acute coronary syndrome, rather than being grouped with other heart conditions, form a small tightly connected cluster with coronary thrombosis (cluster “E”). This shows that clinical connection of these heart conditions to coronary thrombosis is very strong and specific.

Above we demonstrate that close alignment of diseases in the embedding space can result from common or related molecular or physiological mechanisms. This property of disease embeddings can be leveraged for generating hypothesis about novel molecular mechanisms of pathological conditions and for suggesting potential drug targets for their treatment. It also helps to uncover unexpected mechanistic similarities between seemingly unrelated pathologies and to guide drug repositioning based on such connections. These goals are achieved by combining disease embedding with gene-disease association data. First, it produces co-embedding of genes and diseases which allows distilling the most relevant molecular mechanisms underlying disease groups. For example, Fig. 5B shows that gene embeddings reveal common electrophysiological pathways for the cluster of cardiovascular pathologies. For comparison, pathway enrichment of the set of all genes annotated in DisGeNET to diseases in this cluster is dominated by inflammation and signal transduction processes that are much less specific to these conditions (Fig. 5C). Second, combination of disease similarity and gene-disease association data allows generating and ranking hypotheses about novel gene-disease associations by “propagating” evidence across clusters of similar conditions. Importantly, this approach does not require knowledge of any disease-associated genes for the test disease itself, making it applicable for finding genes related to poorly characterized pathologies and orphan diseases if there are data on their co-occurrence with other conditions.

Clustering of diseases in the embedding space plays a crucial role in these applications. Since embedding vector of a gene is the sum of normalized embedding vectors of all its associated diseases, genes that are exclusively associated with a group of related diseases are co-embedded with corresponding disease cluster. At the same time, prolific genes that are associated with a range of diverse conditions do not selectively co-embed with any of the disease types. This is illustrated on Fig 8 which shows how clustering of diseases affects distribution of their associated genes in the embedding space. This effect is reflected in pathway analysis of the cluster of cardiovascular diseases (Fig. 5). Genes related to muscle electrophysiology are very specific to these conditions and are closely aligned with this disease cluster in the embedding space. Therefore, these genes are selected by our algorithm and take part in the pathway enrichment. On the other hand, inflammation processes are very common across the spectrum of different conditions with the result that inflammation-related genes are not systematically co-embedded with cardiovascular diseases and are effectively excluded from functional analysis. Thus, disease clustering drives selection of disease type-specific genes for functional characterization of disease clusters. In the process of generating hypotheses about novel gene-disease associations and drug targets, disease clustering allows leveraging dependencies among pathologies that exist on different physiological levels and may not be immediately evident from known disease mechanisms. For example, we hypothesized a connection between pulmonary embolism and MMP9 based on its association with aortic aneurisms and myocardial infarction, diseases that are clustered with PE in the embedding space (Fig. 7). While MMP9 is not directly involved in the etiology of PE, it plays an important role in vascular remodeling. In addition to aortic aneurisms, this gene mediates other conditions such as heart failure (47) which can arise as a consequence of pulmonary embolism and lead to death in PE patients (48,49). Some studies show that inhibition of MMP9 leads to significant decrease in mortality in the animal model of PE (43-46), making it a promising drug target for this disease. This example illustrates how unexpected mechanistic connections between diseases, revealed by disease embedding, can produce hypotheses about gene-disease associations and can be leveraged to suggest novel drug targets. Overall, this “information propagation” approach yields approximately 276K gene-disease pairs with the aggregated evidence score of > 1 which are not among annotated associations in DisGeNET. They represent a large pool of hypotheses about novel gene-disease associations. These hypotheses are further ranked by the values of their scores and can be mined as a source of drug targets and biomarkers.

**Figure 8.**
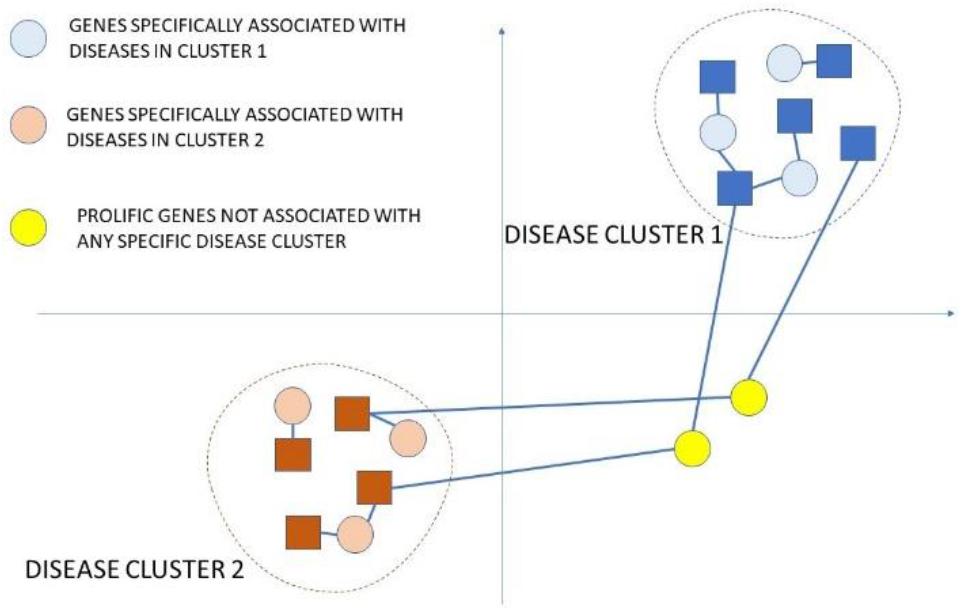
Alignment of genes with disease clusters in the embedding space. Genes that are specific to a set of similar diseases co-cluster with these diseases (blue and orange circles). Genes associated with diseases that are not similar to each other are not aligned with any particular cluster (yellow circles).

## Methods

### Datasets

*Cui2vec* dataset was downloaded from https://rdrr.io/github/ML4LHS/clinspacy/man/dataset_cui2vec_embeddings.html as a table containing embeddings of 109,053 Concept Unique Identifiers (CUI) for UMLS terms. Each row contains 500-dimensional embedding vector for a CUI. Identifiers in this set correspond to human diseases and many other biomedical concepts, such as treatments, metabolites, etc.

Gene-disease association data were downloaded from https://www.disgenet.org/website. Direct matching of CUIs between DisGeNET and *cui2vec* tables yields 3,568 common disease identifiers associated with 8,686 genes. This corresponds to approximately 32% of diseases and 63% of annotations in DisGeNET. Further analysis is limited to this subset of genes and diseases. The disease embedding matrix (***D***) is created using corresponding subset of the *cui2vec* table.

### Disease clustering

Cosine similarity between diseases is defined as 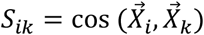, where 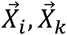 are disease embedding vectors. Distance is defined as *d*_*ik*_ = 1 − *S*_*ik*_. Hierarchical clustering of the resulting 3,568*X*3,568 disease-disease distance matrix is performed using *hclust* function in R with “ward.D2” algorithm (50). Different values of clustering height cut-off are applied to identify sets of disease clusters.

### Disease MeSH mapping

MeSH categories were downloaded from https://www.nlm.nih.gov/databases/download/mesh.html. 26 top MeSH categories are used in the analysis. For each disease cluster a 26-component vector is computed with each component corresponding to the fraction of cluster’s diseases belonging to each one of the top MeSH categories. Note that diseases in MeSH may belong to multiple categories, therefore the sum of fractions across all 26 categories may exceed 1 for some clusters. Pearson correlation is computed between vectors corresponding to different clusters and average value of correlation is computed across all possible cluster pairs. The procedure is repeated for different cluster height cut-off, and average Pearson correlation is computed for all pairs of clusters-MeSH mapping vectors to construct the plot of correlation vs. the number of clusters (Fig. 3).

### Gene-disease co-embedding

First, we create gene-disease association matrix ***A*** with 3,568 rows corresponding to disease and 8,686 columns corresponding to genes. For the purposes of this study, we disregard confidence values provided by DisGeNET. Therefore, the elements of this matrix are “1” if corresponding association is present in DisGeNET and “0” otherwise. Next, we generate a normalized version of the disease embedding matrix 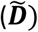 by dividing each row by the length of the corresponding embedding vector. This is done to achieve equal contribution of each gene-associated disease to the gene’s embedding vector, regardless of the lengths of disease embedding vectors. Finally, the gene embedding matrix ***G*** is computed as a product of the transpose of gene-disease association and normalized disease embedding matrices: 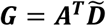.

### Relating genes to disease clusters

To identify genes related to disease clusters, we define embedding vector for each disease cluster as the average direction of normalized embedding vectors of diseases in the cluster. Next, we compute cosine similarity between gene embedding vectors and cluster embedding vectors for each of the disease clusters. By our definition, a gene is associated with the cluster with which it has the greatest value of similarity. Thus, each gene is associated with only one disease cluster – the one with which it is most closely aligned in the embedding space. The procedure is illustrated on Figure 9.

**Figure 9.**
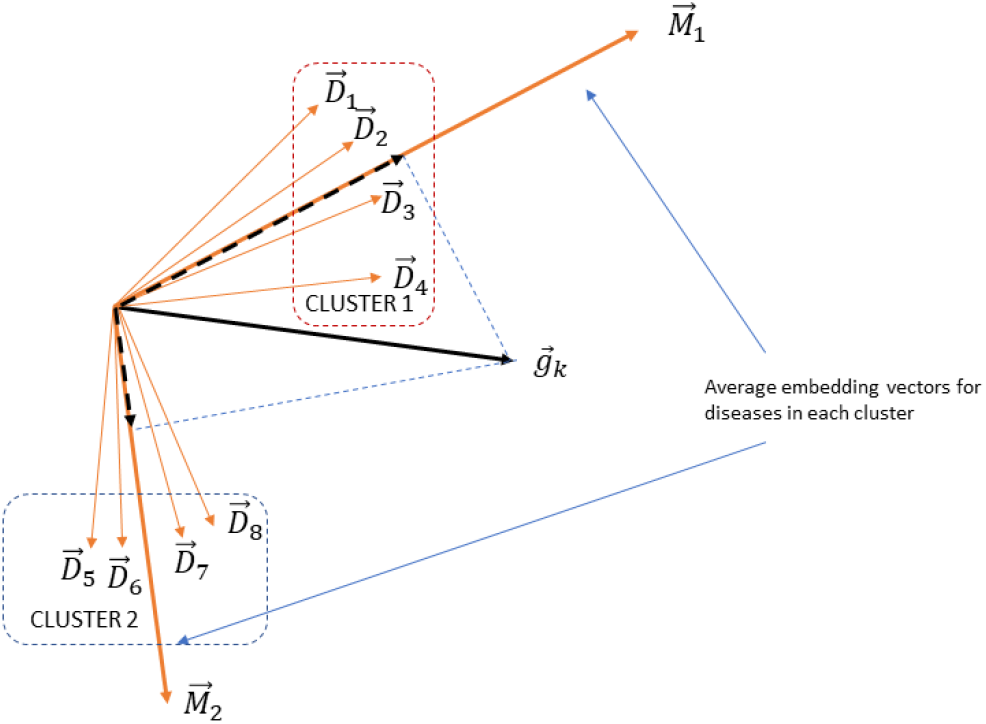
Associating genes with disease clusters to identify genes most closely associated with disease types.

### Scoring putative gene-disease associations

Aggregated evidence scores (*Ã*_*Ã*_) for putative associations between disease *i* and gene *j* are computed by adding up contributions from other diseases associated with gene *j* in proportion to their *transformed similarity* with disease *i*:

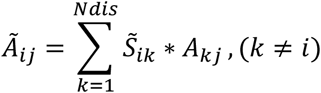

where 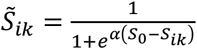 is transformed similarity, 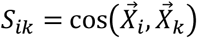, *S*_0_ = 0.8 is a similarity threshold, *α* = 20 is a coefficient which regulates the steepness of the cut-off. Calculation of the aggregated similarity scores is performed by matrix multiplication: 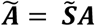.The rationale behind using transformed similarity is our assumption that only sufficiently similar diseases provide meaningful information and should contribute to the aggregated score, whereas contributions from weakly similar diseases mostly represents noise. The choice of the value of the similarity threshold *S*_0_ is based on the following procedure. First, for each disease we find the most similar neighbor and note the value of similarity. Second, we calculate the average of these values across all diseases and round it down to the lower tenth. Therefore, we ensure that for most diseases there will be at least one but no more than a handful of contributing neighbors.

Once aggregated evidence scores are computed we use them as predictors of gene-disease associations and compute areas under the ROC curve (AUC) to evaluate quality of the predictions. We assume that this method may give good predictions for some diseases, while not working well for others. Therefore, we compute AUC separately for each disease. We consider all genes annotated to a disease as *true positives*. While we do not have the knowledge of *true negatives* – genes that are truly not associated with a disease, we assume that gene–disease association matrix is sparse - the majority of not associated genes represent the true lack of associations (rather than the lack of knowledge). Therefore, we form sets of “control” genes for every disease by sampling all genes lacking association records for the corresponding disease. To maintain balanced ratios between cases and controls, the size of the sample in each case is equal to the number of disease-associated genes. Furthermore, random sampling is repeated 100 times for every disease and average value of AUC is calculated over 100 groups of “control” genes.

### Computing numbers of gene-disease similarity pairs

Randomizations are performed with the following procedure:

- 20 random permutations of row labels of disease embedding matrix are performed. This preserves overall clustering of disease vectors in the embedding space, but each cluster is now composed of a set of randomly selected diseases.
- For each permutation, gene embeddings are computed using real gene-disease associations and permutated disease labels.
- Numbers of gene-disease similarity pairs exceeding certain similarity thresholds are counted for a set of similarity thresholds in the range 0.2-0.95.
- Ratios between the number of similarity pairs corresponding to real disease embeddings vs. average number of pairs obtained with permuted disease labels are computed for each of the threshold values.

