## Supplemental tables information for "Identification of disease mechanisms and novel disease genes using clinical concept embeddings learned from massive amounts of biomedical data"

### Supplementary Tables

**TableS1.** Hierarchical clustering of 3,568 diseases based on the cosine similarity of their embedding vectors. Columns A-C: Disease CUI ID; Disease Name, Disease Semantic Type (as provided in DisGeNET). Columns D-M: cluster numbers for each disease at corresponding cluster height cut-offs (h).

**TableS2.** Fractions of cluster's disease annotated to each of the 26 top MeSH categories.
